# Thymic rejuvenation via induced thymic epithelial cells (iTECs) from *FOXN1*-overexpressing fibroblasts to counteract inflammaging

**DOI:** 10.1101/2020.03.17.995357

**Authors:** Jiyoung Oh, Weikan Wang, Rachel Thomas, Dong-Ming Su

## Abstract

Age-associated systemic, chronic, sterile inflammatory condition (inflammaging) is partially attributed to increased self (auto)-reactivity, resulting from disruption of central tolerance in the aged, involuted thymus. Age-related thymic involution causally results from gradually declined expression of the transcription factor forkhead box N1 (*FOXN1*) in thymic epithelial cells (TECs), while exogenous *FOXN1* in TECs can significantly rescue age-related thymic involution. Given the findings that induced TECs (iTECs) from *FOXN1*-overexpressing embryonic fibroblasts can generate an ectopic *de novo* thymus under the kidney capsule and intra-thymically injected natural young TECs can lead to middle-aged thymus regrowth, we sought to expand upon these two findings by applying them as a novel thymic rejuvenation strategy with two types of promoter-driven (*Rosa26*CreER^T^ and *FoxN1*Cre) Cre-mediated iTECs. We engrafted iTECs, rather than natural young TECs, directly into the aged thymus and/or peri-thymus and found a significantly rejuvenated architecture and function in the native aged murine thymus. The engrafted iTECs drove regrowth of the aged thymus in both male and female mice, showing not only increased thymopoiesis, but also reinforcement of thymocyte negative selection, thereby, reducing senescent T cells and auto-reactive T cell-mediated inflammaging phenotypes in old mice. Therefore, this is a promising thymic rejuvenation strategy with preclinical significance, which can potentially rescue declined thymopoiesis and impaired negative selection to significantly, albeit partially, restore the defective central tolerance and reduce subclinical chronic inflammatory symptoms in the elderly.

**Graphical Abstract:** 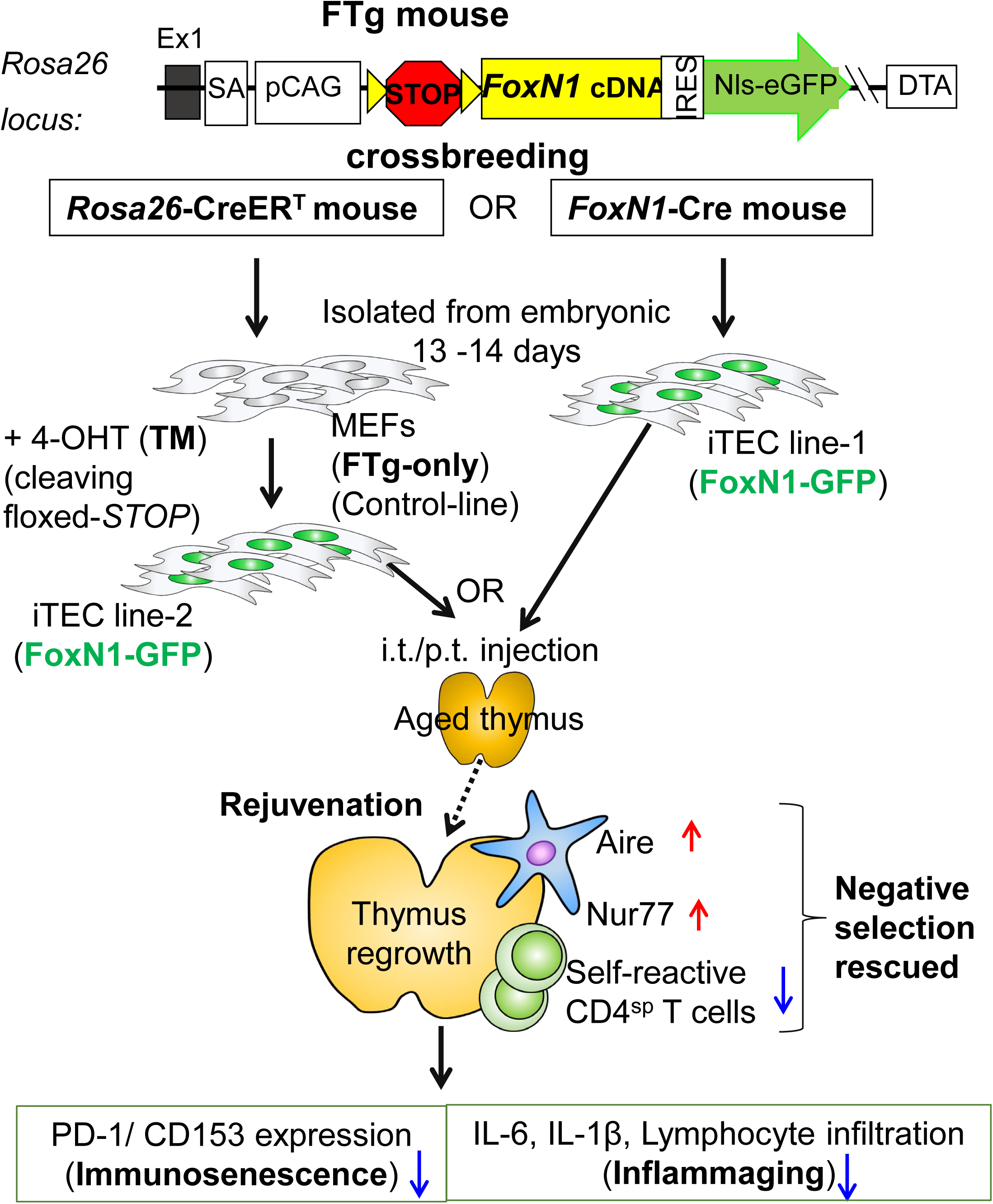

A novel rejuvenation strategy via the *FOXN1*-TEC axis using induced two types of *FOXN1*-overexpressing embryonic fibroblasts (termed iTECs) by intrathymic injection is able to counteract age-related thymic involution, which rescued negative selection, thereby, reducing peripheral T cell-associated inflammaging conditions.

## INTRODUCTION

Age-related immune dysfunction is generally characterized by two extremes: immunosenescence (immune insufficiency) (McElhaney & Effros 2009) and inflammaging (a chronic, persistent, sterile systemic inflammation, partially due to strong self-reactivity) (Freund *et al.* 2010; Franceschi & Campisi 2014). These are antagonistic phenotypes, but they actually comprise two sides of the same coin (Fulop *et al.* 2017), and are associated with functional defects in the aged, atrophied thymus (Goronzy & Weyand 2012; Xia *et al.* 2012; Coder *et al.* 2015; Palmer *et al.* 2018). Immunosenescence, unlike cultured cellular senescence, happens at systemic levels exhibiting diminished immune reaction in response to antigen stimulations, mainly due to contracted T cell receptor (TCR) repertoire diversity (Vallejo 2006). This is primarily attributed to a declined output of naïve T cells from the aged, atrophied thymus (Hale *et al.* 2006) and expansion of monoclonal memory T cells in the periphery (Detailed in our review) (Thomas *et al.* 2020). Although inflammaging was originally attributed to somatic cell senescence-associated secretory phenotype (SASP) (Coppe *et al.* 2010) and chronic innate immune activation (Fulop *et al.* 2017; Fulop *et al.* 2018), the contribution of aged adaptive immune components and specifically self-reactive T lymphocytes, as a probable primary contributor, has been recently determined (Coder *et al.* 2015; Fulop *et al.* 2018). The increased self-reactive T cells in the elderly are derived from perturbed central T cell tolerance establishment (Xia *et al.* 2012; Coder *et al.* 2015; Klein *et al.* 2019), due to defects in negative selection and altered regulatory T (Treg) cell generation (Coder *et al.* 2015; Oh *et al.* 2017) in the aged, atrophied thymus.

During aging the thymus undergoes a progressive, age-related atrophy, or involution, and a key trigger is the primary defect in thymic epithelial cells (TECs), which is mainly attributed to gradually diminished expression of transcription factor forkhead box N1 (*FOXN1*) in TECs (Ortman *et al.* 2002; Sun *et al.* 2010; Rode *et al.* 2015). Therefore, in order to ameliorate immunosenescence and reduce inflammaging through restoration of the aged T cell immune system, many have focused on targeting TECs in the aged thymus. Since the TEC-autonomous factor *FOXN1* is heavily implicated in onset and progression of age-related thymic involution, currently more strategies for rejuvenation of the aged thymic concentrate on the *FOXN1*-TEC axis, although there are strategies other than *FOXN1*-based, such as growth/sex hormones (Detailed in our review) (Thomas *et al.* 2020). *FOXN1*-TEC axis strategies includes *FoxN1*^eGFP/+^ knock-in epithelial cells (Barsanti *et al.* 2017), newborn TEC-based intrathymic injection (Kim *et al.* 2015), inducible *FoxN1*-expressed mouse embryonic fibroblast (MEF)-based ectopic thymus generation (Bredenkamp *et al.* 2014b), and genetically-based rejuvenation via enhancement of exogenous *FoxN1* expression with *FoxN1* cDNA plasmid (Sun *et al.* 2010) and *FoxN1* transgene (Tg) in TECs (Zook *et al.* 2011; Bredenkamp *et al.* 2014a). In addition, cytokine/growth factor-to-TEC based rejuvenation strategies have been studied, including addition of mesenchymal cell-derived keratinocyte growth factor (KGF) (Min *et al.* 2007), macrophage- and T lymphocyte-derived insulin-like growth factor-1 (IGF-1) (Chu *et al.* 2008), thymic stromal cell-derived bone morphogenetic protein-4 (BMP4) (Tsai *et al.* 2003; Wertheimer *et al.* 2018), and lymphoid tissue inducer (LTi) cell-derived IL-22 (Dudakov *et al.* 2012). These factors are produced from cells of mesenchymal or hematopoietic origin, but target non-hematopoietic TECs associated with up-regulating *FoxN1* expression in TECs. Finally, epigenetically-based rejuvenation, via extracellular vesicles and exosomes extracted from young healthy serum has been shown to rejuvenate not only the peripheral T cell system, but also the thymus by enhancing *FoxN1* expression (Wang *et al.* 2018). Therefore, there is potential for rejuvenating thymic aging by primarily targeting the restoration of TEC homeostasis through rescuing age-related declined *FoxN1* expression.

Among the *FOXN1*-TEC axis therapies for thymic rejuvenation, two strategies are particularly attractive. One strategy is to aggregate induced *Rosa26(R26)*CreER^T^-mediated *FOXN1*-overexpressed MEFs (converting these cells into pseudo-TECs, termed induced TECs or iTECs) along with early-stage thymocytes and fetal mesenchymal cells to build an ectopic thymus under the kidney capsule of adult mice (Bredenkamp *et al.* 2014b). This *de novo* ectopic thymus produced functional T cells. However, one limitation is that the aged, native thymus remains in the host releasing self-reactive T cells that still contribute to inflammaging. The other strategy is an intrathymic injection of freshly isolated newborn TECs (non-manipulated TECs), in which *FoxN1* is normally highly expressed, into the native thymus of middle-aged mice (Kim *et al.* 2015). This led to restoration of thymopoiesis. However, collection of fresh newborn TECs is not feasible when considering translating this rejuvenation strategy to humans, and isolation of fresh TECs without thymocyte contamination is very difficult since TECs and their progenitors comprise a miniscule portion of the thymus (Ulyanchenko *et al.* 2016). Therefore, these promising thymic rejuvenation strategies for development of a practical therapy contain several limitations.

Fortunately, fibroblasts, which could be very easily isolated from human patients, can be engineered to overexpress *FOXN1* for induction of iTECs for intra-/peri-thymic injection. Based on these scientific premises, we expanded on these two findings and applied them to develop a novel thymic rejuvenation strategy. We directly engrafted iTECs into the aged, native thymus to rejuvenate function of the aged, native thymus and assessed this in a mouse model, by using MEFs from our engineered *STOP*^flox^-*FoxN1* transgenic (FTg) mouse allele (Zhang *et al.* 2012; Ruan *et al.* 2014) (Supplemental Fig. S1), mediated by two types of promoter-driven (*Rosa26*CreER^T^ and *FoxN1*Cre) Cre-recombinase.

We found that the engrafted iTECs drove regrowth of the aged thymuses in both male and female mice with increased thymopoiesis and improved thymic architecture. These led to a reinforcement of thymocyte negative selection in the native, aged thymus, thereby attenuating auto-reactive T cell-mediated inflammaging phenotypes and reducing senescent T cells in old mice. Although the native, aged thymus cannot fully return to young levels in our system, this is a promising thymic rejuvenation strategy with preclinical significance to counteract inflammaging.

## RESULTS

### Preparation and characterization of iTECs

A previous report demonstrated that enforced *FOXN1* expression in MEFs from embryos generated by crossbreeding of *STOP*^flox^-*FoxN1* transgenic and *R26-CreER*^T^ mice induced epithelial characteristics in fibroblasts (Bredenkamp *et al.* 2014b). Since we generated similar *STOP*^flox^-*FoxN1* transgenic (exogenous *FoxN1* cDNA driven by *R26* promoter, termed FTg) mice (DNA construct is shown in Supplemental Fig. S1) (Zhang *et al.* 2012; Ruan *et al.* 2014), we crossbred these mice with either *R26-CreER*^T^ or *FoxN1-Cre* mice to generate FTg:*R26CreER*^T^ and FTg:*FoxN1Cre* embryonic mice, respectively. We confirmed epithelial characteristics in MEFs from two different promoter-driven *FoxN1* expressing lines in our mouse colonies (Fig. 1). Using NIs-eGFP (nuclear localization signal enhanced green fluorescent protein) as an indicator of exogenous *FoxN1* expression (Zhang *et al.* 2012; Ruan *et al.* 2014) in the cultured MEFs (isolated from embryonic day-13 (E13) and E14 mice), we found MEFs from FTg-only (without any Cre-Tg) and FTg:*R26CreER*^T^ without addition of tamoxifen (xTM, 4-OHT) did not express GFP (Fig. 1A left panels) due to lack of activated Cre, while FTg:*FoxN1Cre* (TM not required) and FTg:*R26CreER*^T^ lines treated with TM for 48 hours showed GFP expression (Fig. 1A right panels and Fig. 1B middle and right panels). We also found that MEFs with greatly increased exogenous *FoxN1* expression from FTg:*FoxN1Cre* and FTg:*R26CreER*^T^ (xTM) mice showed TEC identifying markers (EpCAM^+^ and MHC-II^+^ cells in the GFP^+^ population) (Fig. 1B, middle and right panels), but not MEFs of FTg:*R26CreER*^T^ without addition of TM (Fig. 1B, left panels).

**Figure 1.**
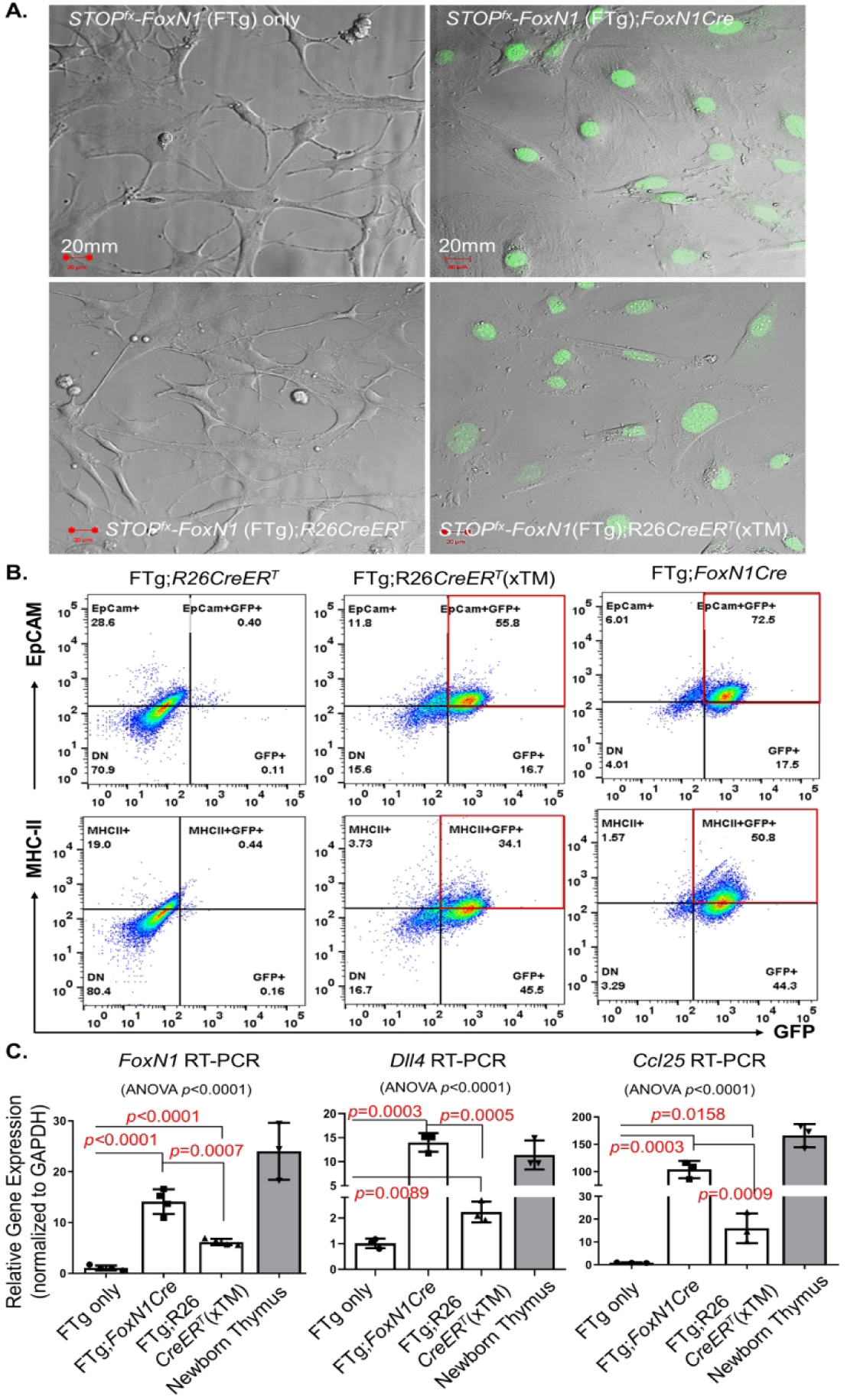
Preparation and characterization of MEFs and iTECs. Mouse embryonic fibroblasts (MEFs) were isolated via trypsinized digestion from E13 and E14 embryonic mice, and cultured in plates with or without 4-hydroxy tamoxifen (symbol: xTM). **(A).** Representative live images from confocal microscopy show MEFs expressed GFP, which represents exogenous FoxN1 (right panels) and was driven by either endogenous *FOXN1*-carried Cre-recombinase at 3’UTR (FTg:*FoxN1Cre*; top-right) or *R26*-carried CreER^T^ treated with TM (FTg:*R26CreER*^T^ xTM; bottom-right); and panels without GFP (left panels) due to either no Cre transgene or no active Cre; **(B)** Representative flow cytometric dot plots (EpCAM vs. GFP: top panels; and MHC-II vs. GFP: bottom panels), in which MEFs expressing GFP (FTg:*R26CreER*^T^ xTM and FTg:*FoxN1Cre*) are termed iTECs (in red boxes of the middle and right panels), compared to MEFs that did not express GFP (FTg:*R26CreER*^T^ due to Cre inactivated without TM treatment – left panels); **(C)** Summarized gene (*FoxN1, Dll4*, and *Ccl25*) expression (via RT-PCR) in cells of four groups: (1) FTg-only: without Cre; (2) FTg:*FoxN1Cre*: Cre expression was endogenously turned on in *FoxN1*^*+*^ cells; (3) FTg:*R26CreER*^T^ xTM: Cre was activated via TM induction, and (4) a newborn thymus control group. A Student *t*-test was used to determine statistical significance and *P* values are shown between every two groups. In addition, an ordinary one-way ANOVA *p*-value summary by comparing multiple groups is shown on top of each panel. All *p*-values were calculated by mean ± SD and “n” animal numbers. Scales showed in each bar are SEMs. Each symbol represents cells from an individual embryonic sample.

Exogenous *FoxN1* mRNAs were indeed increased in the two Cre-activated groups (Fig. 1C, two middle bars in leftmost panel). In addition, some TEC functional molecules, which are key effectors in promoting thymocyte development, such as Notch ligand *Dll4* and thymus-expressed chemokine ligand *Ccl25*, were increased in MEFs with activated *Cre*-Tg (Fig. 1C, middle and right panels). Notably, expression of both exogenous *FoxN1* and effector molecules were increased in the iTECs, but their increased levels in these pseudo-TECs were still lower or similar to their expression in the natural newborn thymus, during which these molecules should be normally highly expressed (rightmost striped bars in Fig. 1C all panels). In addition, *FoxN1*Cre-mediated expression of exogenous *FoxN1* and effector molecules in the FTg:*FoxN1Cre* line was higher than *R26*CreER^T^-mediated ones. This is probably due to *Cre-*Tg turning on via endogenous *FoxN1 in vivo*, which is activated by E11.25 in the thymus (Gordon *et al.* 2001) and potentially in the E12.5 skin (Gordon *et al.* 2007) or alternatively at low levels in the E13.5 skin (Bredenkamp *et al.* 2014b) during the organogenesis of B6 mice. This *in vivo* endogenous *FoxN1*-induced exogenous *FoxN1* expression is 48hrs earlier than *in vitro* TM-induced expression in the FTg:*R26CreER*^T^ line. Together, Cre-induced expression of exogenous *FoxN1* and TEC functional molecules in MEFs conferred TEC characteristics to these MEFs. Therefore, these MEFs were termed as iTECs.

### Intra-/peri-thymic (i.t./p.t.) transplantation of iTECs drove aged thymus regrowth

Previous reports demonstrated that intrathymic (i.t.) injection of fetal thymic cells, containing young TECs with high-levels of *FoxN1* expression, into middle-aged (9-12 months old) mice drove recipient thymus growth and increased T cell production (Kim *et al.* 2015). Since Kim *et al*.*’s* approach requires a newborn thymus for rescuing an aged thymus and newborn TECs are difficult to obtain and purify, we tested whether our iTECs could yield similar outcomes in fully aged (over 18 months old) mice. We firstly examined thymus regrowth and thymopoiesis of aged mice (∼18 months old at the time of the injection and 19 - 20 months old at the time of analysis) after i.t./p.t. transplantation of iTECs. Our results show that transplantation of iTECs indeed drove aged thymus regrowth (Fig. 2) exhibited by increased thymic size, weight, and thymocyte numbers (Figs. 2A, 2B, and 2C, respectively). These changes were the same in male and female mice. Although these improvements did not reach the same levels as the young mice (Fig. 2A, top row and Figs. 2B and C, leftmost group in each panel), it was significantly improved compared to the naturally aged group without transplantation of iTECs (transplantation of FTg-only MEFs served as a negative control allowing for the same surgical stress as the iTEC-engrafted groups).

**Figure 2.**
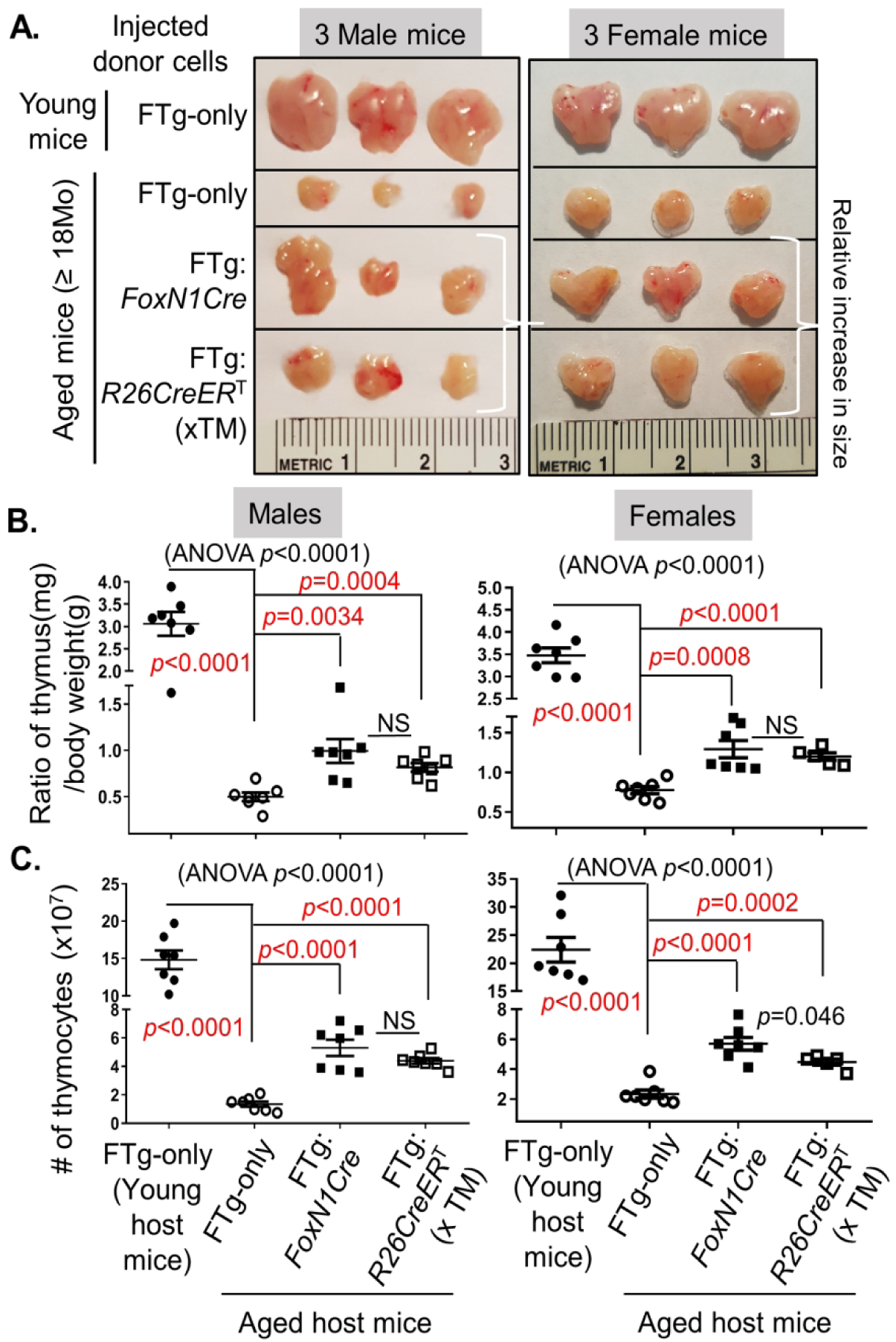
Transplantation of iTECs drove re-growth of the aged thymus in both male and female mice. Naturally aged mice (WT, ≥18 months old at cellular transplantation; 20 – 21 months old at analysis) were intra-/peri-thymically (i.t./p.t.) transplanted with FTg-only MEFs or either of two promoter-driven exogenous *FoxN1* expressing iTECs; one group of young mice served as a control. Forty-five days after engraftment, the thymic mass was analyzed. **(A)** Representative images of the thymuses engrafted with donor cells; **(B)** Ratios of thymus/body weight and **(C)** Results of absolute thymocyte numbers per thymus from donor cell-engrafted aged male and female mice (one young group, leftmost, served as control); Statistical analysis, data expression, and each symbol per animal are the same as Fig. 1.

Overall, our iTECs better resemble newborn TECs and more efficiently drive the aged (≥18 months old), atrophied thymus regrowth and rejuvenation of thymopoiesis. It appears that the efficacy from both iTEC lines were generally similar, but endogenous *FoxN1* promoter-driven Cre was slightly better than *R26* promoter-driven CreER^T^ (xTM) (See Fig. 2C rightmost two groups in the rightmost panel). This could be explained by the fact that although *R26*CreER^T^ is turned on *in vitro* during the culture with TM-induction, which is 48 hours later than the *FoxN1*Cre is activated *in vivo*. Expression of the effector molecules in FTg:*R26*CreER^T^ line was lower than FTg:*FoxN1*Cre line (Fig. 1C), but they could become the equivalent after injection into the host thymuses, since the effector molecules probably increase only to a homeostatic plateau.

### Grafted iTECs rejuvenated thymic architecture in aged mice

Increased thymic mass (Figs. 2A and B) generally reflects expansion in thymocytes (Fig. 2C) and regrowth in TECs, because rejuvenation of TEC meshwork is essential for thymocyte regrowth. We examined TEC-based thymic microstructure using TEC-associated markers (Fig. 3). After co-staining with keratin-5 (K5, red) (medullary region) and K8, (cortical region) (all in green in Fig. 3), the aged, atrophied thymus showed disorganized and reduced K5^+^ regions (Fig. 3A, the second panel from left). After treatment with either FTg:*FoxN1Cre* or FTg:*R26CreER*^T^ (xTM) iTECs (Fig. 3A, right 2 panels), the K5^+^ regions became organized, similar to the young thymus (Fig. 3A, leftmost panel). Increased UEA-1^+^ TECs showed the well-organized medulla, exhibiting the same trends as the K5^+^ region (Fig. 3B). Claudin (Cld)-3 and -4 (Cld3+4) are immature medullary thymic epithelial cells (mTEC) markers (Hamazaki *et al.* 2007; Sekai *et al.* 2014) and β5t is mainly expressed in immature cortical thymic epithelial cells (cTECs) (Ripen *et al.* 2011). These were decreased in the naturally aged thymus, but were rescued in the naturally aged thymus treated with either of the two promoter-driven Cre-induced iTECs (Figs. 3C and D). These results infer that input of iTECs enhances native TEC regrowth to rejuvenate aged thymic architecture, thereby improving thymic microenvironment and rebooting thymopoiesis.

**Figure 3.**
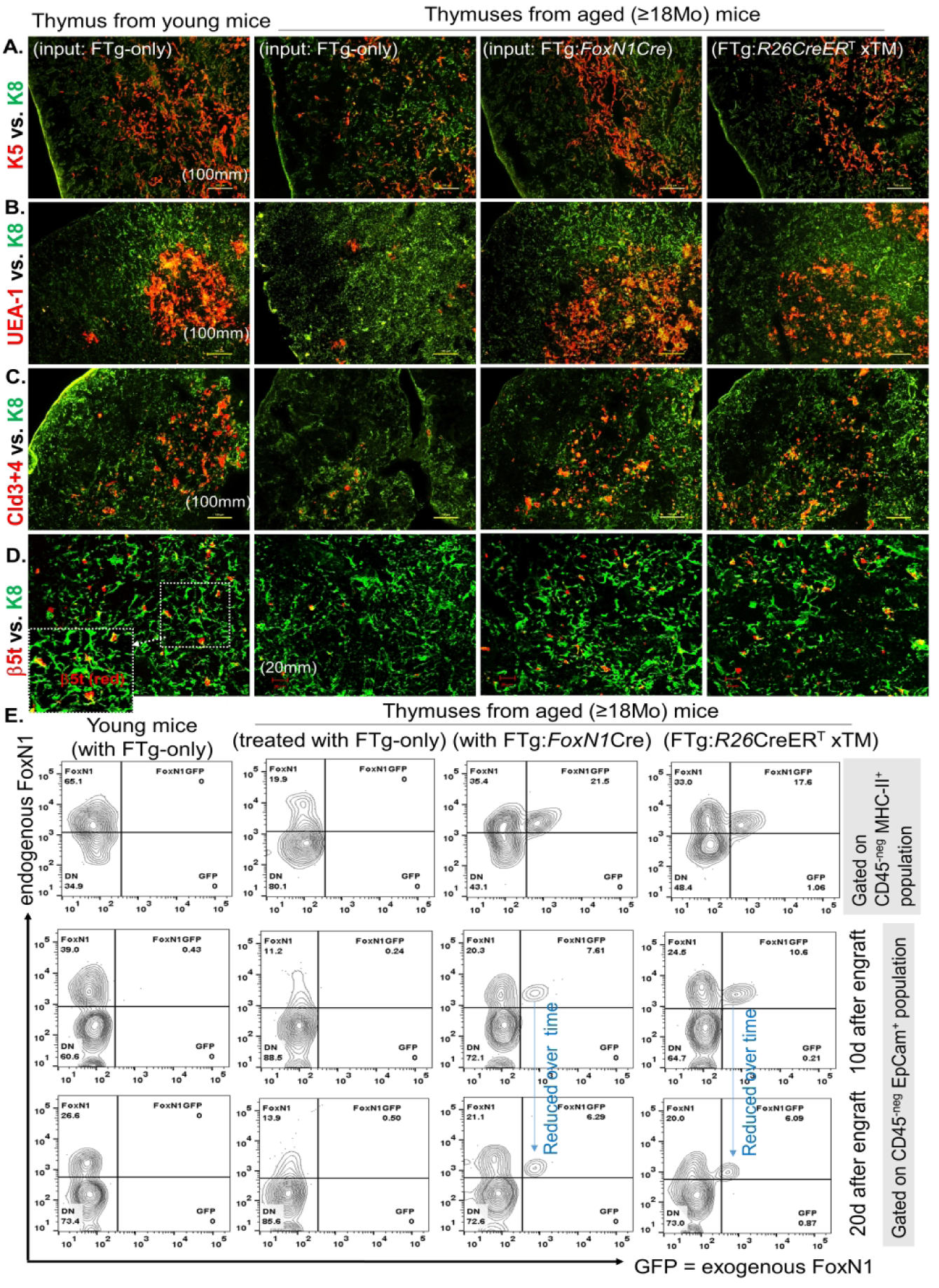
Transplantation of iTECs rejuvenated thymic architecture of aged mice via both exogenous iTEC growth and endogenous TEC regrowth. Same experimental setting as described in Fig. 2. Cryosections of the thymic tissue (Representative immunofluorescence images shown in panels A –D) were co-stained with various immunofluorescence antibodies for TEC developmental and architectural profiles. **(A)** K5 (red) vs. K8 (green); **(B)** UEA-1 (red) vs. K8 (green); **(C)** Claudin (Cld)-3+4 (red) vs. K8 (green); **(D)** β5t (red) vs. K8 (green). Data are representative of 3 biological replicates in each group with essentially identical results. (**E**) Flow cytometric analysis of endogenous TECs (FoxN1^+^GFP^-neg^) and exogenous TECs (from iTECs, FoxN1^+^GFP^+^) in the mTECs (CD45^-^MHC-II^+^) or pan-TECs (CD45^-^EpCam^+^) of various thymuses, 10 or 20 days after engraftment with MEFs or iTECs, based on endogenous FoxN1 (by antibody) and exogenous FoxN1 (by GFP) expression.

To confirm whether the observed TECs regrew from the native aged thymus when they received stimulation from iTEC-rejuvenated microenvironment, or if these TECs grew directly from newly transplanted iTECs, we examined the sources of these TECs in the rejuvenated, aged thymuses based on endogenous and exogenous FoxN1 expression. The TECs with positive staining for FoxN1 using rabbit anti-FoxN1 (the antibody was kindly provided by Dr. Itoi, Japan) (Itoi *et al.* 2007) exhibited only endogenous FoxN1, while the TECs with both antibody-specific FoxN1 staining and FTg-GFP (See supplemental Fig.S1) expression (double positive) contain exogenous FoxN1 and would therefore be derived from the newly transplanted iTECs. We found that both native TECs and transplanted iTECs were expanding within 10 days after the engraftment (Fig. 3E, right two-ranked panels on top and middle two rows), particularly in mTECs (CD45^-neg^MHC-II^+^ population, top row). Further, the transplanted iTECs exhibited reduced expansion but the native TECs were still robustly expanding over 20 days after the engraftment (Fig. 3E bottom row). The results suggest that although engrafted iTECs growth is transient, they do exhibit growth, and they can also promote native TEC growth in the recipient thymus even after their growth begins to wane. Thus, it seems that once native TECs receive necessary stimulation, they undergo a more prolonged expansion compared to the engrafted iTECs. However, both the engrafted iTECs and rejuvenated native TECs cooperate to restore the aged thymic microenvironment to promote thymocyte expansion.

### Engrafted iTECs expanded *Aire*-expressing mTECs, increased negative selection signaling in CD4^SP^ thymocytes, and restored declined thymocyte negative selection in the aged thymus

Autoimmune regulatory, *Aire*, gene is expressed by mTECs to mediate self-antigen expression and promote central immune tolerance via thymocyte negative selection and Treg generation (Anderson *et al.* 2005; Anderson & Su 2016). In the aged thymus Aire-expressing mTECs are disrupted and/or declined (Coder *et al.* 2015; Wang *et al.* 2018). Since transplantation of iTECs enhanced biological characteristics of native TECs in the naturally aged thymus (Fig. 3), we tested whether transplantation of the two iTEC lines was able to expand declined Aire-expressing mTECs and found positive results (Fig. 4A. bottom row) with statistical significance (Fig. 4B. two right groups) in the aged thymus.

**Figure 4.**
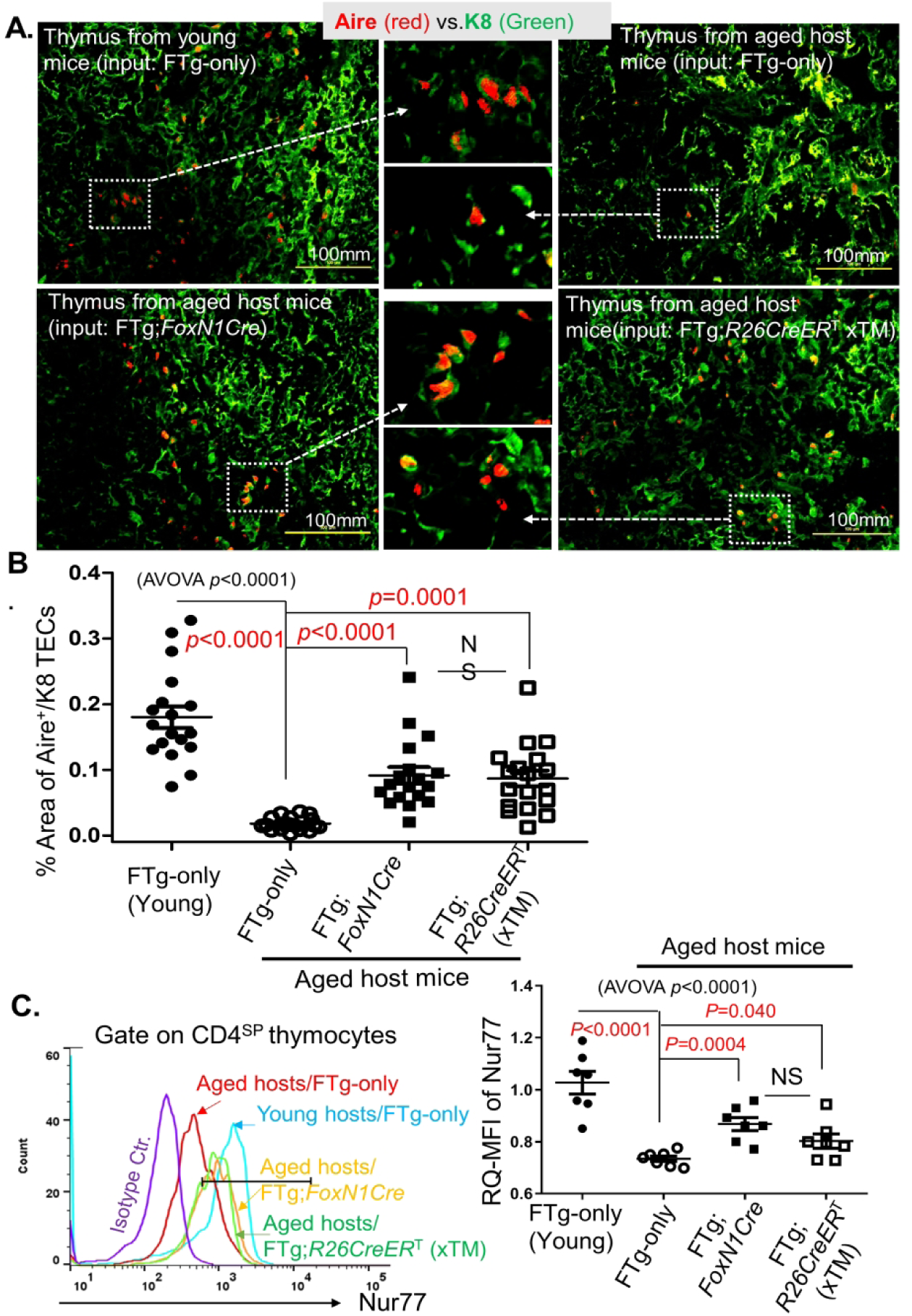
Transplantation of iTECs boosted *Aire* gene expression in the age thymus and showed enhanced negative selection signaling strength via Nur77 in CD4^SP^ thymocytes of aged mice. Same experimental setting as described in Fig. 2. **(A)** Representative immunofluorescence staining images of Aire^+^ TECs (red) in K8^+^ TEC counterstaining (green). Data are representative of three biological replicates in each group with essentially identical results; **(B)** Summarized result shows the percent area of Aire^+^ TECs against K8^+^ counterstaining based on the slides in panel A. Each symbol represents one thymic tissue section; ±6 thymic tissue sections at disconnected locations (non-sequential slides) from an individual mouse thymus were counted with Image-J software; **(C)** Flow cytometric results show increased Nur77 signaling strength [relative quantitative (RQ) mean fluorescent intensity (MFI)] in CD4^SP^ thymocytes of young (control) or aged mice that were engrafted with MEFs or two types of iTECs. Left panel: histogram of Nur77 MFI in CD4^SP^ thymocytes; Right panel: Nur77 RQ-MFI in CD4^SP^ populations of various groups. Statistical analysis, data expression, and each symbol per animal are the same as Fig. 1.

Self (auto)-reactive thymocytes undergo negative selection dependent on TCR signaling strength, while the intensity of Nur77 expression in thymocytes reflects a negative selection signaling strength. We examined mean fluorescence intensity (MFI) of Nur77 in CD4 single positive (CD4^SP^) thymocytes from various groups (Fig. 4C), and found MFIs of Nur77 in CD4^SP^ thymocytes were indeed increased in the two iTEC-grafted groups (Fig. 4C, right two square-symbol groups in the right panel). Although these increases did not reach the same levels as in young mice (Fig. 4C, a filled-circle group in the right panel), they were significantly increased, compared to naturally-aged controls (FTg-only group).

The results provided an indication that transplantation of iTECs potentially restores TEC function in negative selection as demonstrated by increased Aire^+^ mTECs and enhanced negative selection signaling strength in the CD4^SP^ thymocytes in the aged thymus. In order to obtain direct evidence that the declined thymocyte negative selection in the aged thymus is really restored, we designed an observable negative selection model, in which mOVA-Tg host young and aged mice were reconstituted with donor OT-II TCR-Tg mouse bone marrow (BM) cells. This is a well-designed thymocyte negative selection model, in which a neo-self-antigen mOVA presented on mTECs induces OT-II TCR-Tg CD4^SP^ thymocyte depletion (negatively selected), able to be observed through flow cytometry assay (Hubert *et al.* 2011; Coder *et al.* 2015). The thymuses in the immune system-reconstituted young and aged mice were engrafted with FTg:*FoxN1*Cre iTECs or control FTg-only MEFs. Four weeks after the transplantation of these cells, the proportion of OT-II-specific CD4^SP^ thymocytes was determined (Fig. 5A). Increased proportion of OT-II-specific CD4^SP^ thymocytes in the mOVA-Tg thymic microenvironment means defective negative selection, which was seen in the aged, atrophied thymus (Fig. 5B middle panels, and Fig. 5C middle bar). However, this proportion was reduced after transplantation with iTECs (Fig. 5B right panels and Fig. 5C rightmost bar) in the aged mOVA-Tg thymuses. Meanwhile, signaling of negative selection (Nur77) in the specific CD4^SP^ thymocytes was increased (Figs. 5D yellow histogram and 5E rightmost bar). The results imply that engrafted iTECs were indeed able to significantly restore mTEC-mediated function for self-reactive thymocyte negative selection in the aged, atrophied thymus.

**Figure 5.**
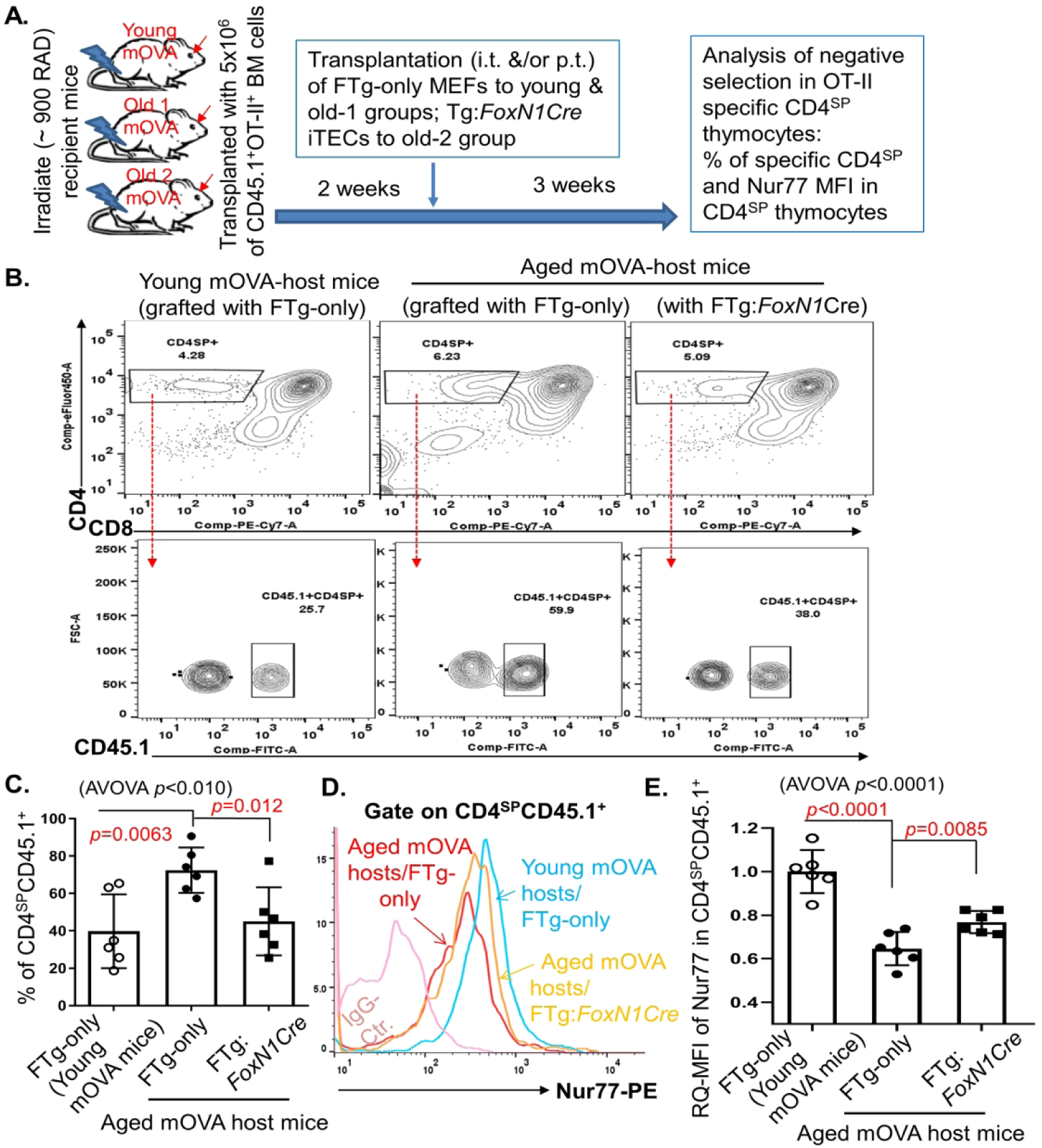
Transplantation of iTECs partially rescued declined thymocyte negative selection in aged mice. **(A)** Reconstituted mOVA-Tg aged mice (mOVA-Tg young mice for control) with OT-II TCR-Tg bone marrow (BM) cells via a ∼900Rad irradiation were intra-/peri-thymically transplanted with MEFs (FTg-only) or iTECs (FTg:*FoxN1Cre*). Negative selection of OT-II TCR-Tg specific CD4^SP^ (CD4^+^CD8^-neg^) thymocytes in the host mOVA-Tg TEC microenvironment was analyzed with a flow cytometer. (**B)** Flow cytometric gating scheme of CD4 vs CD8 (top row) and engrafted donor BM (CD45.1^+^) produced OT-II specific TCR-Tg CD4^SP^ thymocytes (bottom row). **(C)** Summarized results of % OT-II specific TCR-Tg CD4^SP^ thymocytes. **(D)** A representative histogram of Nur77 MFI in OT-II specific TCR-Tg CD4^SP^ thymocytes. **(E)** Relative quantitative (RQ)-mean fluorescent intensity (MFI) of Nur77 signaling strength in OT-II specific TCR-Tg CD4^SP^ thymocytes, by setting RQ-MFI in young thymocytes as 1.0 (i.e. signaling with 100% intensity). Statistical analysis, data expression, and each symbol per animal are the same as Fig. 1.

### Engrafted iTECs counteracted inflammaging by exhibiting reduced inflammatory cytokines and lymphocyte infiltration into non-lymphoid organs in the periphery

To confirm whether the restoration of negative selection in the iTEC-engrafted aged thymus could counteract inflammaging-associated phenotypes in the aged periphery, we examined the levels of inflammatory cytokines and lymphocyte infiltration into non-lymphoid organs through adoptive transfer of rejuvenated spleen cells from rejuvenated mice. As we know, inflammaging is attributed to not only senescence somatic cells producing SASP and chronic innate immune cell activation, but also self (auto)-reactive T cell-induced self-tissue damages. These self-reactive T cells are released from the aged, atrophied thymus due to defective negative selection (Goronzy & Weyand 2012; Xia *et al.* 2012; Coder *et al.* 2015; Palmer *et al.* 2018). If the engrafted iTECs can restore declined negative selection, the self-reactive T cells released should be reduced, and thereby, peripheral inflammaging-associated phenotypes should be attenuated.

We examined two types of inflammatory phenotypes. One is the levels of two classic pro-inflammatory cytokines (IL-6 and IL-1β) in the serum of the naturally aged mice, 45 days after engraftment with iTECs or control MEFs. As reported, these cytokines were increased in the serum of naturally-aged mice (Fig. 6A, the opened diamond-symbol group), but they were significantly decreased after engraftment with either type of iTECs (Fig. 6A, the opened and filled square-symbol groups).

**Figure 6.**
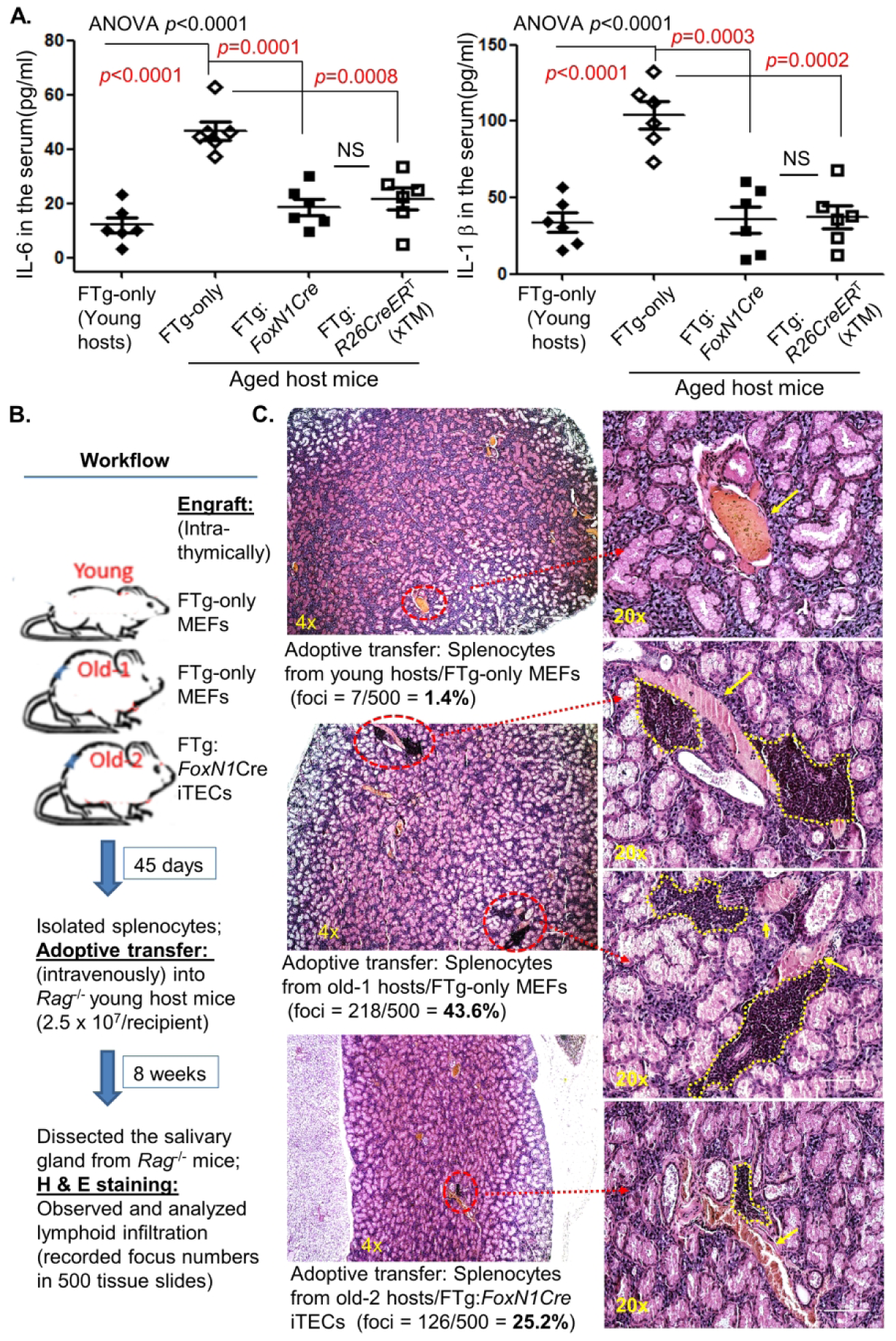
Transplantation of iTECs attenuated inflammaging-associated phenotypes by reducing inflammatory cytokines and lymphoid cell infiltration into non-lymphoid organs in aged mice. **(A)** Serum was collected from mice with the same treatment as in Fig. 2. Concentration of pro-inflammatory cytokines IL-6 (Left panel) and IL-1β (Right panel) in pg/mg of serum protein was measured through an ELISA method. Statistical analysis, data expression, and each symbol per animal are the same as Fig. 1. **(B)** Workflow of adoptive transfer, showing that splenocytes (2.5 x 10^7^ cells per recipient mouse) from rejuvenated and control young or aged WT mice were transferred via i.v. injection into young *Rag*^-/-^ recipient mice. Eight weeks after the transfer, the salivary glands were subjected to analysis of lymphocyte infiltration; **(C)** Representative H&E stained images of the salivary glands from the adoptive transfer *Rag*^-/-^ recipient mice, showing foci of lymphoid cell infiltration (red circles in 4x images and yellow circles in 20x images). Data are representative of 500 tissue slides from 3 animals in each group, and numbers of infiltration foci in 500 tissue slides and the % of lymphoid cell infiltrated foci are shown.

The second phenotype we assessed was lymphocyte infiltration into non-lymphoid tissue (the salivary gland). The approach was the same as described in our previous reports (Coder *et al.* 2015; Wang *et al.* 2018), and the workflow is shown in Fig. 6B. Splenocytes from thymic-rejuvenated mice or control mice were adoptively transferred into lymphocyte-free *Rag*^-/-^ young mice, and lymphocyte infiltration in the salivary gland (Fig. 6C) was observed. We obtained consistent results with inflammatory cytokines, (Fig. 6A) that iTECs were able to reduce lymphocyte infiltration into non-lymphoid salivary gland (Fig. 6C, the bottom panels). The results indicate that engraftment of iTECs into the aged thymus rejuvenated thymic function, which in turn attenuated inflammaging-associated inflammatory phenotypes in aged individuals.

### Engrafted iTECs indirectly reduced senescent T cells and enhanced T cell immune response in the periphery of aged mice

Inflammaging is also partially attributed to immunosenescence because senescent/ exhausted peripheral T cells not only produce inflammatory factors but are also unable to properly clear senescent somatic cells, which produce SASP (Prata *et al.* 2019; Thomas *et al.* 2020). We asked whether iTEC-driven rejuvenation of aged thymic function could counteract inflammaging through reducing senescent T cells associated with increased output of newly-generated T cells, since the rejuvenated thymus increases thymopoiesis (Fig. 2). We found that 45 days after iTEC engraftment senescent CD4^SP^ T cells (CD4^+^PD-1^+^CD153^+^) (Shimatani *et al.* 2009; Tahir *et al.* 2015) were significantly reduced in the periphery of aged mice (Supportive Figs. S2A and B, right two panels and right two bars), compared to the aged mice which received FTg-only MEFs.

In addition, we also verified the peripheral CD4^SP^ T cell response to co-stimulation from CD3ε and CD28 antibodies. This response, represented by intracellular IL-2 mean fluorescence intensity (MFI) (Fig. S2C), was declined in peripheral CD4^SP^ T cells of aged individuals (Fig. S2D, the 2^nd^ bar from the left) (Sun *et al.* 2010), but was significantly restored in peripheral CD4^SP^ T cells from iTEC-rejuvenated mice (Fig. S2D, two bars with filled and opened square symbols), implying increased proportion of newly-generated T cells in the rejuvenated mice. Taken together, iTEC-driven changes in the aged thymus could additionally confer a positive rejuvenation effect on the peripheral T cell system.

## DISCUSSION

T cell-mediated adaptive immunity during aging is intricately involved in both immunosenescence and inflammaging. One of the potential strategies for ameliorating these two extremes is rejuvenation of the aged, involuted thymus. Restoring thymic function of central tolerance establishment via repairing the defects in negative selection is critical for counteracting inflammaging. Although there are many strategies for rejuvenation of thymic involution, targeting defective TEC homeostasis via the *FOXN1*-TEC axis is one of the most effective strategies.

We tested an application of cellular rejuvenation of age-related thymic involution by using iTECs generated from *R26*CreER^T^ and *FoxN1*Cre-induced exogenous *FoxN1* in *STOP*^flox^-*FoxN1*-Tg embryonic fibroblasts with intrathymic injection. We found that the engrafted iTECs were able to induce aged thymus regrowth with increased thymopoiesis in aged male and female mice (Fig. 2), in which native TECs were reorganized (Figs. 3A-D, right two columns) and underwent expansion (Fig. 3E, right two columns). We also observed reinforced thymocyte negative selection (Figs. 4 and 5). This resulted in reduced auto-reactive T cell-mediated inflammaging-associated phenotypes and diminished peripheral senescent T cells in the aged periphery (Figs. 6 and S2).

The underlying mechanism of thymic rejuvenation potentially involves restoration of TEC regrowth in the aged thymus via both expanded engrafted iTECs (increased GFP^+^FoxN1^+^, double positive, TECs) and induced expansion of native TECs (increased GFP^-neg^FoxN1^+^, single positive, TECs) (Fig. 3E, right two columns). This improves the aged thymic microenvironment, promoting normal thymocyte homeostasis and development. These effects culminated in attenuation of inflammaging phenotypes (Fig. 6) and removal of senescent T cells (Figs. S2A and S2B). Although we did not directly measure native T cells (T cells generated prior to iTEC engraftment), we found that T cells from rejuvenated mice exhibited an increased response to TCR stimulation (Figs. S2C and S2D), which is a functional sign of healthy newly-generated T cells.

Although the rejuvenation was partial, since it cannot restore to the same levels as in young mice, it was significant when compared to the same aged counterparts treated with non-exogenous *FoxN1*-expressing MEFs. The effects of a one-time transplantation of these cells is also most likely transient, since the engrafted iTECs are not TEC stem cells and therefore do not demonstrate unlimited growth after engraftment in the aged, native thymus. Compared to the generation of an ectopic *de novo* thymus with induced *FOXN1*-overexpressing MEFs under the kidney capsule of adult mice (Bredenkamp *et al.* 2014b) and intrathymic injection of newborn TECs into the middle-aged thymus (Kim *et al.* 2015), our strategy facilitates a more clinically translational rejuvenation therapy. Although an ectopic *de novo* thymus can generate naïve T cells, this does not remedy the increased self-reactive T cells released by the native atrophied thymus remaining in the aged host. In addition, intrathymic injection of newborn TECs can rejuvenate middle-aged thymus in mice (Kim *et al.* 2015), but the source of newborn TECs for human treatment is limited. Further, our rejuvenation effects were observed in aged mice (≥18 months old) rather than limited to middle-aged mice (Kim *et al.* 2015).

In comparison with exogenous *FoxN1* expression and rejuvenation effects from two promoter-driven *Cre*-Tg (*FoxN1Cre* and *R26CreER*^T^)-mediated iTECs, exogenous *FoxN1* expression was slightly higher in the former cell type (Fig. 1C leftmost panel), and the effects were not that different between the two lines (Figs. 2-4). We think that this is probably due to the length of time for which the exogenous *FoxN1*-Tg has been activated. It has been turned on *in vivo* before their isolation, because endogenous *FoxN1*-driven-Cre could have been already activated in the E13 and E14 MEF cells, whereas, the exogenous *FoxN1*-Tg expression mediated by *R26*CreER^T^ is turned on after dissection and during the 48-hour culture with TM induction, i.e. 48 hours later than the former. However, the effector molecules (*Dll4* and *Ccl25*) most likely reach a homeostatic plateau. Once these two lines are injected into host mice, expression of the effector molecules in the *R26*CreER^T^ line could feasibly “catch up” to the levels expressed by the former line. In addition, *FoxN1Cre* mediates exogenous *FoxN1* expression only in skin epithelial cells of MEFs, while *R26CreER*^T^ mediates exogenous *FoxN1* expression in most tissues, including fibroblasts and epithelial cells of MEFs, since the *R26* promoter is ubiquitous. Thus, it is not surprising that the effects from both lines are similar. The results imply that this cellular therapeutic strategy is highly clinically translational, since fibroblasts derived directly from patients themselves, who would be treatment recipients, can be readily targeted for genetic engineering of *FoxN1* expression.

In sum, our preliminary, proof-of-principle, cellular-based rejuvenation strategy via the *FOXN1*-TEC axis with intra-/peri-thymic injection is a promising thymic rejuvenation strategy with potential clinical significance. Once the application study is further formulated and investigated, intrathymic transplantation of genetically engineered *FoxN1-*expressing patient skin cells (fibroblasts) could facilitate attenuation of T cell immunosenescence and subclinical chronic inflammatory symptoms in the elderly.

## Experimental Procedures

### Animal models

C57BL/6 genetic background mouse models were used. Wild-type (WT) young and aged mice were from our breeding colonies and National Institute on Aging (NIA) aged rodent colonies. *STOP*^flox^-*FoxN1* transgenic (FTg) mice were generated in our lab previously (Zhang *et al.* 2012; Ruan *et al.* 2014) (Supportive Figure S1) and were crossbred with either *R26*-*CreER*^T^ mice (Jackson Lab #004847) or *FoxN1*-*Cre* mice (Jackson Lab #018448) for the generation of FTg:*R26CreER*^T^ [tamoxifen (TM)-inducible exogenous *FoxN1* overexpression in the *R26*-expressing tissues] and FTg:*FoxN1Cre* [exogenous *FoxN1* overexpression induced by endogenous *FoxN1* promoter-driven Cre-Tg (Gordon *et al.* 2007)] embryonic mice, respectively. Other genetically engineered mouse colonies were RIP-mOVA [the rat insulin promoter (RIP)-driven membrane-bound ovalbumin] Tg mice (Jackson Lab #005431); OT-II^+^ TCR-Tg (transgenic TCR recognizing ovalbumin in the context of MHC-class II, I-A^b^) mice (Jackson Lab #004194); and *Rag*^-/-^ (*Rag1* gene knockout) mice (Jackson Lab #002216). Mouse ages are indicated in each figure legend, or defined as young (1 - 2 months old) and naturally-aged (±18 months old). All animal experiments were performed in compliance with protocols approved by the Institutional Animal Care and Use Committee of the University of North Texas Health Science Center, following guidelines of the National Institutes of Health.

### Preparation of MEFs for intrathymic injection

MEFs were prepared from E13 and 14 embryonic mice (the gestation day-0 “E0” was determined by the presence of a vaginal plug in the first morning on the mother mouse). All the organs of the embryonic mice were removed except for the body with skin, which was trypsinized with Trypsin-EDTA solution to generate single-cell suspensions. Cells were cultured in 10%FBS/DMEM medium, with 2mM L-glutamine, 1mM pyruvate and 50μM 2-mercaptoethanol. In cultured E13 and 14 embryonic FTg:*FoxN1Cre* MEFs (i.e. one type of iTEC donor cells), exogenous *FoxN1* is consistently expressed, due to endogenous *FoxN1*-driven Cre having been turned on, which was found at part of the skin of E12.5 embryonic mice (Gordon *et al.* 2007), and spontaneously activated at low levels, which was observed at E13.5 MEFs (Bredenkamp *et al.* 2014b). For inducing exogenous *FoxN1* overexpression in FTg:*R26CreER*^T^ MEFs (i.e. another type of iTEC donor cells), 1μM of 4-hydroxy tamoxifen (TM) (4-OHT) was added in the culture for 48hr. Exogenous *FoxN1* overexpression [based on green fluorescent protein (GFP) expression] in the two types of MEF lines was examined after 48hr culture. These two cell lines were expanded with two passages. We used the third passage cells as injection reagents. FTg-only (without any Cre Tg) MEFs (control donor cells) were used as negative control. All embryonic mice for preparation of MEFs were genotyped. All cells were checked for GFP expression prior to engraftment.

### Intra-/peri-thymic (i.t./p.t.) injection of donor cells into recipient mice

FTg-only MEFs (negative control) and two types of promoter-driven Cre-mediated FTg iTECs were injected at 1×10^6^ cells in 20μl of volume per recipient mouse (young or naturally aged) into the thymus and/or peri-thymus in three locations via a suprasternal notch surgery under anesthesia (Burnley *et al.* 2013). Forty-five days after the injection, the tissues of the recipient mice were analyzed for various phenotypes. More details about the operation are depicted in **Supportive Experimental Procedures**.

### Bone marrow (BM) adoptive transfer for assessing negative selection

Erythrocyte-depleted and mature T cell-depleted (via anti-CD3 MACS beads and columns, Miltenyi Biotech) BM cells from OT-II^+^ TCR-Tg mice, which carry a copy of CD45.1 congenic marker, were intravenously (i.v.) injected into recipient young or aged mOVA-Tg mice at 5 × 10^6^ cells per recipient mouse, which had received irradiation at doses of ∼900 Rad. Two weeks after the BM cell transfer, FTg-only MEFs and FTg:*FoxN1Cre* iTECs were intrathymically (i.t.) injected into the thymus/peri-thymus of the recipient mOVA-Tg mice. Four weeks after the engraftment, the thymuses of the recipient mOVA-Tg mice were dissected for analysis of negative selection (proportion of CD4^SP^ and MFI of Nur77 in CD4^SP^).

### Transplantation of splenocytes into Rag^-/-^ recipients for assessing lymphocyte infiltration

Protocol per our previous publication (Coder *et al.* 2015): briefly, erythrocyte-depleted splenocytes from FTg-only MEF- or FTg:*FoxN1Cre* iTEC-engrafted young or aged WT mice were i.v. injected at 2.5 × 10^7^ cells per recipient mouse into the young recipient *Rag* ^-/-^ mice. Eight weeks after the transplantation, the salivary glands from the young recipient *Rag* ^-/-^ mice were analyzed for lymphocyte inflammatory infiltration with Hematoxylin and Eosin (H&E) staining in paraffin sections (5μm thick).

### General analysis methods

Detailed analysis methods (Real-time RT-PCR, flow cytometer, immunofluorescence staining, and ELISA, etc.), as well as reagents are described in **Supportive Experimental Procedures**.

### Statistics

Either the unpaired two-tailed Student’s *t*-test for comparing two groups with equal variance or one-way ANOVA with Bonferroni correction for comparing multiple groups were employed. Differences were considered statistically significant at values of * *p* < 0.05; ** *p* < 0.01; *** *p* < 0.001. All statistics were analyzed with Prism-8 software (GraphPad).

## Abbreviations

Aire: autoimmune regulator gene;
BM: bone marrow;
CD4^SP^: CD4^+^CD8^-neg^ single positive;
CreER^T^: Cre-recombinase and estrogen-receptor fusion protein;
cTEC/mTEC: cortical/medullary thymic epithelial cells;
FTg: *STOP*^flox^-*FoxN1*cDNA transgene;
GFP: green fluorescent protein;
iTECs: induced TECs from inducible *FOXN1*-overexpressing embryonic fibroblasts;
MEF: mouse embryonic fibroblast;
mOVA: the membrane-bound chicken ovalbumin driven by the rat insulin promoter (RIP);
OT-II CD4^+^ T cells: MHC class-II restricted and OVA recognizing T cell receptor transgenic CD4^+^ T cells;
*R26*: *Rosa26* gene;
*Rag*: *V(D)J*-recombination-activating gene;
SASP: senescence-associated secretory phenotype;
SP: single positive;
TCR: T cell receptor;
Tg: transgenic;
TM: tamoxifen;
WT: wild type.

## Author contribution

J.O. designed and performed the most experiments, analyzed data, prepared figures, and wrote the manuscript; W.W. performed most of the hands-on animal work; R.T. performed part of the experiments, helped to write and proofread the manuscript; D-M.S. conceived, designed, and supervised the project, helped with hands-on animal work, analyzed data, and wrote the manuscript.

## Author declaration

All authors have no conflicts of financial interests associated with this manuscript.

