## Supplemental Methods & Figures for "Thymic rejuvenation via induced thymic epithelial cells (iTECs) from *FOXN1*-overexpressing fibroblasts to counteract inflammaging"

### Supportive Experimental Procedures

#### *Real-time RT-PCR for gene expression*

Total RNAs were isolated from FTg-only MEF cells or two types of iTEC cells using TRIzol reagent (Invitrogen), then reversed transcribed (RT) into cDNA with the SuperScriptIII cDNA kit (Invitrogen/ThermoFisher Scientific). Real-time RT-PCR was performed with TaqMan reagents and primers of *FoxN1* (#Mm00433948), *Dll4* (Mm00444619), *Ccl25* (Mm00436443), and house-keeping gene, GAPDH, from Thermo Fisher scientific. The relative difference between samples were calculated using $\Delta\Delta C_T$ methods, GAPDH was used to normalize samples for comparison, as previously described (Oh *et al.* 2017).

#### *Flow cytometry assays of single suspended cells*

Single cells suspensions [homogenized thymocytes or splenocytes, or Collagenase-V/DNase-I dissociated TECs (Burnley *et al.* 2013; Sizova *et al.* 2018)] were stained with various fluorochrome-conjugated antibodies (from BioLegend unless otherwise indicated) and analyzed with an LSR-II flow cytometer (BD Biosciences) and FlowJo software (Oh *et al.* 2017). For combinative cell surface and intracellular staining, cells were stained with multiple fluorochrome-conjugated antibodies on cell surface, blocked with an Fc receptor antibody (CD16/CD32), and followed by fixation/permeabilization solution (eBioscience, Cat. #88-8824-00), then intracellular staining, such as Nur77, IL-2, etc. (Oh *et al.* 2017).

#### *Tissue sectioning with immunofluorescence*

Dissected mouse tissues were prepared as 6μm-thick cryo-sections, and stained with various primary antibodies and fluorochrome-conjugated secondary antibodies as described previously (Burnley *et al.* 2013; Ruan *et al.* 2014; Sizova *et al.* 2018; Wang *et al.* 2018). The results were visualized by a fluorescence microscope (Nikon Eclipse Ti-U) or confocal laser scanning microscope (LSM 510 Meta, Zeiss), and results were quantified by Image-J software.

#### *ELISA assay for pro-inflammatory cytokines*

Mouse serum was isolated from iTEC cell-engrafted mice, and pro-inflammatory cytokines IL-6 and IL-1β were quantified by ELISA kit (BioLegend Cat. #431304 and #432605) (Sizova *et al.* 2018). Each serum sample was diluted 1:2 with PBS and samples were prepared in duplicate.

IL-6 and IL-1β standard concentration curves were observed within a range of 0-200 pg/ml. Absorbance was measured at 450nm with a BioTek ELx800 ELISA reader.

#### *Image analyses*

Open the immunofluorescent staining overlaid image files (showing both red and green channels) in NIH Image-J software and split this merged double-channel immunofluorescent image into two single-channel images, then convert both images into binary images by clicking “Process”. Click “Analyze” to measure positively stained cells in each layer in percentage (%). Finally, the ratios of % positive areas (in our case, Aire positive staining) versus % counterstaining area (in our case, K8 positive staining) were calculated. One of the representative samples is shown in below.

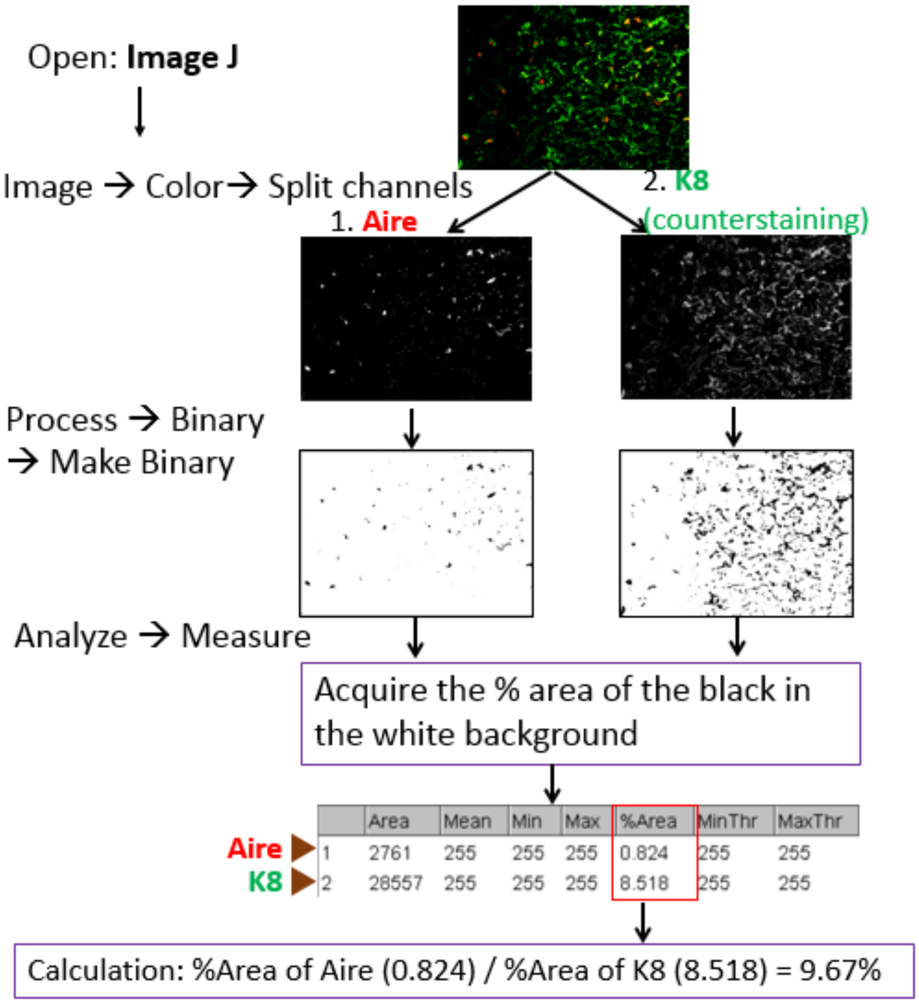

#### *Operation of intra-/peri-thymic injection*

Intra/peri-thymic injection is a routine technique in our Lab (Burnley *et al.* 2013). Anesthetized mouse was laid ventral side up and a sagittal ±0.5cm incision was cut with surgical scissors in the skin on the suprasternal fossa (below Figure-a). A pole was laid under the neck to allow for hyperextension of head. The muscle covered on the suprasternal fossa was moved (below Figure-b, green arrow) with forceps to expose the suprasternal fossa (below Figure-b, red arrow). A Hamilton syringe was injected into the chest through the suprasternal fossa at almost $0^o$ angle along the body toward the tail. Two more injections were made by slightly shifting the needle angles toward the left and right of the body. The total volume injected per adult mouse was 25µl roughly equally distributed in the three differently angled injections. Lab members have practiced this operation many times with injection of trypan blue dye to perfect the accuracy of the injection in both normal thymus (below Figure-c left side) and atrophied thymus (below Figure-c right side).

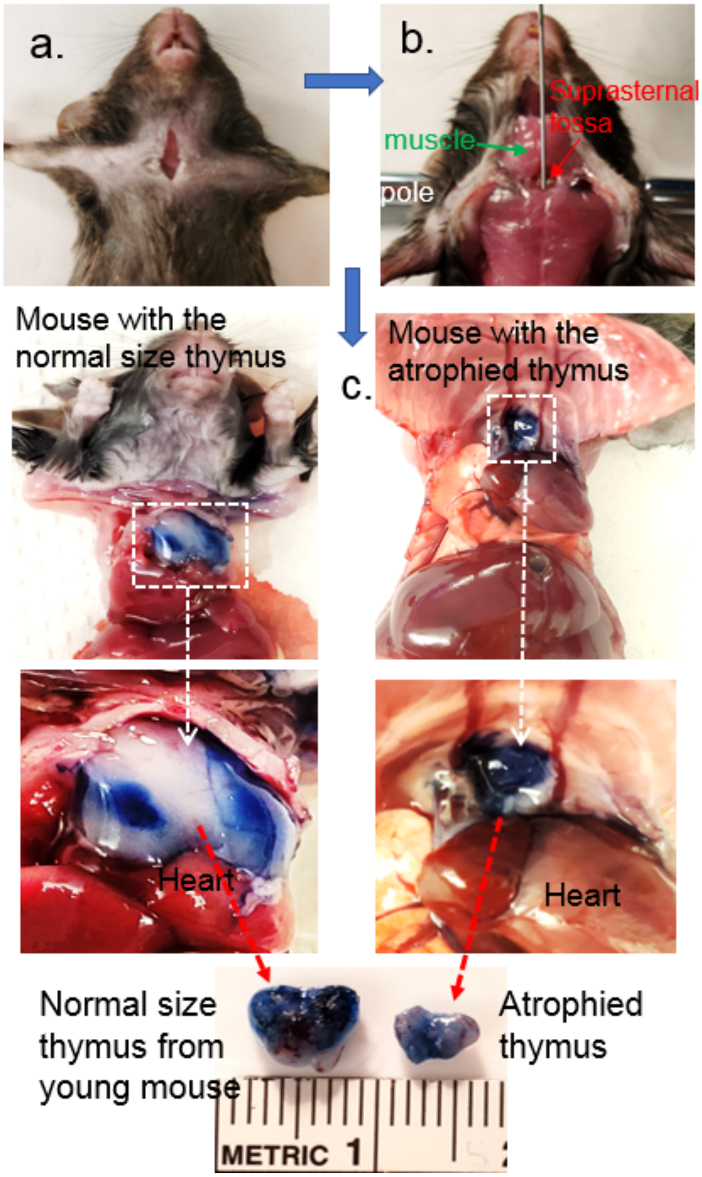

### Supportive Figure Legends

**Fig. S1.** The generation of *R26-STOP*$^{flox}$-*FoxN1*$^{Tg}$ mice. Schematic diagram of *R26-STOP*$^{flox}$-*FoxN1*$^{Tg}$ gene targeting and elements of DNA insertion. Top line: Wild-type *R26* locus; bottom line: insertional DNA in the target vector.

**SA**: splicing acceptor

**pCAG**: poly **C**MV enhancer element, chicken beta-**A**ctin promoter, and rabbit beta-**G**lobin splice acceptor site

**STOP**: DNA sequence cassette for transcriptional stop

**Yellow triangles**: two ***loxp***s

**IRES**: internal ribosome entry site

**Nls-eGFP**: nuclear localization signal (Nls) enhanced green fluorescent protein (eGFP).

**Fig, S2. Transplantation of iTECs reduced senescent T cells and enhanced TCR response in the periphery of aged mice** Same experimental setting as described in Fig. 2. Erythrocyte-depleted splenocytes were either directly stained for senescent T cells or isolated for culture (2x10$^6$ per well) with co-stimulation of anti-CD3ε and anti-CD28 (2μg/ml each) supplemented with GolgiSTOP (0.7μl/ml, BD Biosciences) for 5hr. **(A)** Flow cytometric gating scheme of senescent CD4$^{SP}$ splenic T cells: PD-1$^+$CD153$^+$ (red boxes) in CD4$^{SP}$ population. **(B)** Summarized results of reduced senescent CD4$^{SP}$ T cells after transplantation with either of two promoter-driven Cre-induced iTECs (right two bars), compared to aged mice treated with FTg-only MEFs. **(C)** Flow cytometric gating scheme of CD4$^{SP}$ splenic T cell responses to the TCR co-stimulation of anti-CD3ε and anti-CD28: intracellular IL-2 (APC) levels in CD4$^{SP}$ T cell population. **(D)** Summarized results of increased intracellular IL-2 levels in CD4$^{SP}$ T cells of aged mice after transplanted with either type of iTECs (two bars with filled or opened square symbols), compared to aged mice without iTEC-treatment (FTg-only). RQ-MFI is a relatively quantitative mean fluorescent intensity, in which IL-2 MFI in young CD4$^{SP}$ T cells (leftmost bar) was set as 1.0. (i.e. full responses to the co-stimulation). A Student *t*-test was used to determine statistical significance between two groups, and *P* values are shown between two groups in each panel. In addition, an ordinary one-way ANOVA *p*-value summary by comparing multiple groups is shown on top of each panel. All *p*-values were calculated by mean ± SD and "n" animal numbers. Scales showed in each bar are SEMs. Each symbol represents an individual animal sample.

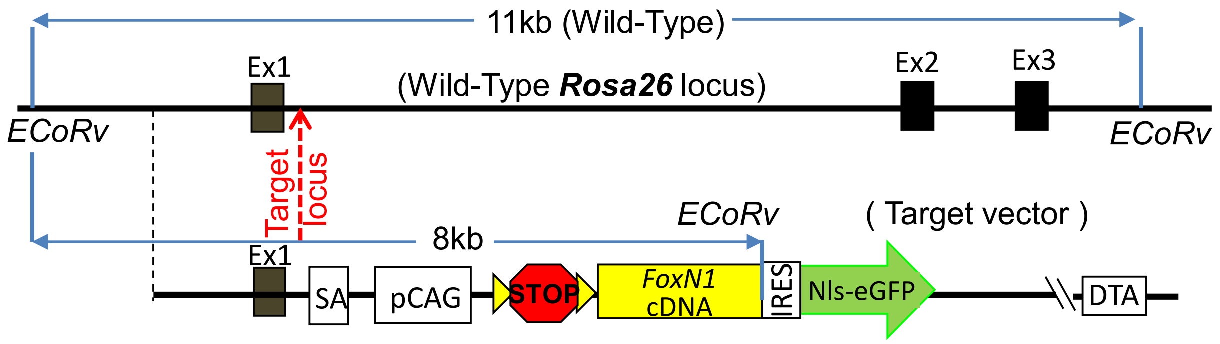


**Fig. S1.** The generation of *Rosa26-STOP*$^{flox}$-*FoxN1*$^{Tg}$ mice. Schematic diagram of *Rosa26-STOP*$^{flox}$-*FoxN1*$^{Tg}$ gene targeting and elements of DNA insertion. Top line: Wild-type *Rosa26* locus; bottom line: insertional DNA in the target vector.

**SA**: splicing acceptor
**pCAG**: poly **C**MV enhancer element, chicken beta-**A**ctin promoter, and rabbit beta-**G**lobin splice acceptor site
**STOP**: DNA sequence cassette for transcriptional stop
**Yellow triangles**: two ***loxp***s
**IRES**: internal ribosome entry site
**Nls-eGFP**: nuclear localization signal (Nls) enhanced green fluorescent protein (eGFP).

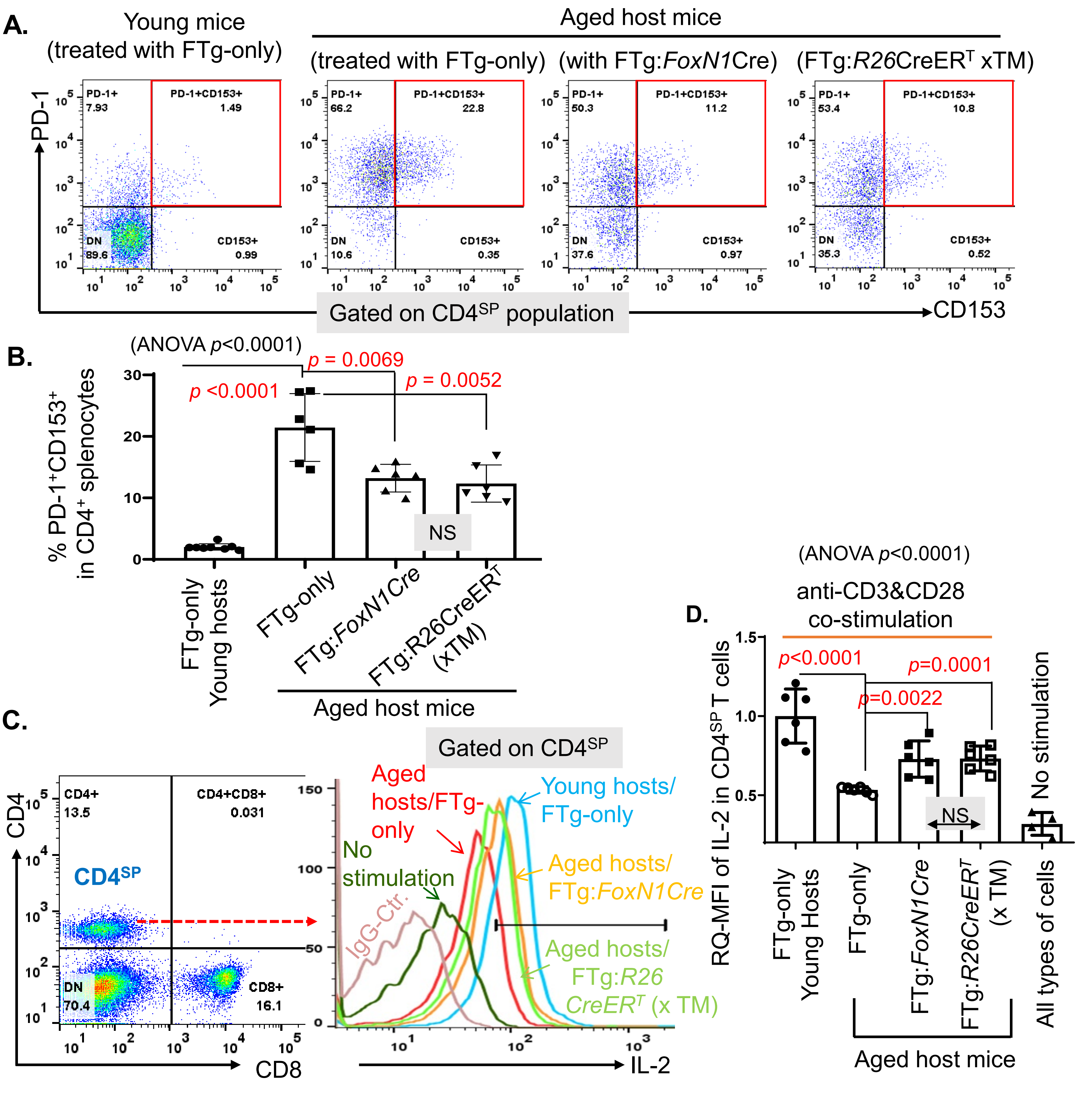
A.
Young mice (treated with FTg-only)
Aged host mice
(treated with FTg-only)
(with FTg:FoxN1Cre)
(FTg:R26CreERT xTM)
PD-1
PD-1+ 7.93
PD-1+CD153+ 1.49
DN 89.6
CD153+ 0.99
PD-1+ 66.2
PD-1+CD153+ 22.8
DN 10.6
CD153+ 0.35
PD-1+ 50.3
PD-1+CD153+ 11.2
DN 37.6
CD153+ 0.97
PD-1+ 53.4
PD-1+CD153+ 10.8
DN 35.3
CD153+ 0.52
Gated on CD4SP population
CD153
B.
(ANOVA p<0.0001)
% PD-1+CD153+ in CD4+ splenocytes
p <0.0001
p = 0.0069
p = 0.0052
NS
FTg-only Young hosts
FTg-only
FTg:FoxN1Cre
FTg:R26CreERT (xTM)
Aged host mice
C.
Gated on CD4SP
CD4
CD4+ 13.5
CD4+CD8+ 0.031
CD4SP
DN 70.4
CD8+ 16.1
CD8
Aged hosts/FTg-only
No stimulation
IgG-Ctr.
Young hosts/ FTg-only
Aged hosts/ FTg:FoxN1Cre
Aged hosts/ FTg:R26 CreERT (x TM)
IL-2
D.
(ANOVA p<0.0001)
anti-CD3&CD28 co-stimulation
RQ-MFI of IL-2 in CD4SP T cells
p<0.0001
p=0.0022
p=0.0001
NS
No stimulation
FTg-only Young Hosts
FTg-only
FTg:FoxN1Cre
FTg:R26CreERT (x TM)
All types of cells
Aged host mice

Fig. S2
